# Measuring the perception and metacognition of time

**DOI:** 10.1101/2023.04.18.537334

**Authors:** Simon J Cropper, Daniel R Little, Liheng Xu, Aurelio M Bruno, Alan Johnston

**Affiliations:** Melbourne School of Psychological Sciences, University of Melbourne, Australia, 3010; Department of Psychology, University of Nottingham, UK; Department of Psychology, University of York, UK

**Keywords:** Time perception, metacognition, ideal Observer Analysis, psychophysical methods

## Abstract

The ability of humans to identify and reproduce short time intervals (in the region of a second) may be affected by many factors ranging from the gender of the individual observer, through the attentional state, to the precise spatiotemporal structure of the stimulus. The relative roles of these very different factors are a challenge to describe and define; several methodological approaches have been used to achieve this to varying degrees of success. Here we describe a new paradigm affording not only a first-order measurement of the perceived duration of an interval but also a second-order metacognitive judgement of perceived time. This approach, we argue, expands the form of the data generally collected in duration-judgements and allows more detailed comparison of psychophysical behaviour to the underlying theory. We also describe a measurement model which provides estimates of the variability of the temporal estimates and the metacognitive judgments allowing comparison to an ideal observer. We fit the model to data collected for judgements of 750ms (bisecting 1500ms) and 1500ms (bisecting 3000ms) intervals across three stimulus modalities (Visual, Audio & Audiovisual). This enhanced form of data on a given interval judgement and the ability to track its progression on a trial-by-trial basis offers a way of looking at the different roles that subject-based, task-based and stimulus-based factors have on the perception of time.

## Introduction

The perception of time from milliseconds to years is a cognitive capacity that fascinates from many different perspectives. Time, as a perceptual quantity, has some unique characteristics which also make it particularly challenging to measure and model. Of all the measurable properties of the world that we do somehow encode and experience, time is arguably the most distanced from its physical realisation when considered in terms of its neural representation. Time is also unique as a perceptual experience in that it can be derived from any sensory modality and much of the work in time perception to date has taken a somewhat modality-agnostic approach, in that visual and auditory sensory inputs have been used approximately equally to code the interval in an experimental setting. Furthermore, the internal awareness of time-passed, the metacognition of time, is critical for the maintenance of an ongoing awareness of, and involvement in, the external world and the segmentation of ongoing external events (Wahlheim et al., 2022; Zacks, 2020; Zacks et al., 2022). The loss of this (temporal) context is a major symptom of psychosis and serious affects the ability of an individual to interact socially or otherwise with their environment (Bocker et al., 2000; Cohen & Dochert, 2005). Time and its passing can be thought of as a thread through the ongoing narrative around which we construct our conscious lives, and while we all have our own ‘thread’ we also rely on substantial agreement about the passing of time to co-exist functionally in the same reality.

The ability of subjects to identify and (re)produce brief temporal intervals is influenced by many factors whether stimulus, subject or task-based (Bruno et al., 2012; Corcoran et al., 2018; Jazayeri & Shadlen, 2010; Piras & Coull, 2011). Several reviews have comprehensively outlined the current state of empirical and theoretical development in time perception (Block & Grondin, 2014; Bruno & Cicchini, 2016; Grondin, 2008, 2010; Matthews, 2015; Matthews & Meck, 2014; Matthews & Meck, 2016; Meck & Ivry, 2016; Tsao et al., 2022); the aim here is not to repeat this information but to introduce and motivate what we believe is a novel and useful method to approach the study of brief to moderate temporal intervals and the metacognition of that perceptual experience. We have used this approach to look at population differences in time perception and metacognition (Corcoran et al., 2018) but examine the methodology and data in a small-N psychophysical design in more detail here, so we may develop our model of timing and metacognition with a higher level of precision. To this end, we examine temporal bisection performance for subsecond (750ms) and suprasecond (1500ms) tasks within visual, audio and audiovisual modalities and the ongoing metacognition of that performance.

### Ways of measuring time

Time is unique as a sensory quantity in that it is not bound to one particular sensory input; from an experimental perspective, the two most common modalities chosen to generate the temporal stimulus are auditory and visual, but measurement of somatosensory time perception is also not uncommon (Tomassini et al., 2011; Watanabe et al., 2010). Grondin (2008; 2010) outlines four main methods of investigating time that have been traditionally distinguished within the literature: verbal-estimation, reproduction, production and comparison. The final ‘comparison’ category can be split into several subcategories (e.g., roving or reminder, single-stimulus, bisection), but this category overall is equivalent to standard psychophysical methods of measurement for any sensory capacity and tends to be favoured in the experimental literature. Time perception from a sensory perspective is generally focused on the examination of durations up to a few seconds. These methods and approaches are neatly summarized in his Figure 1 (Grondin, 2010). The novel method examined in detail here falls under this broad ‘comparison’ definition, and incorporates characteristics of both reproduction and bisection, with an additional metacognitive component.

**Figure 1.**
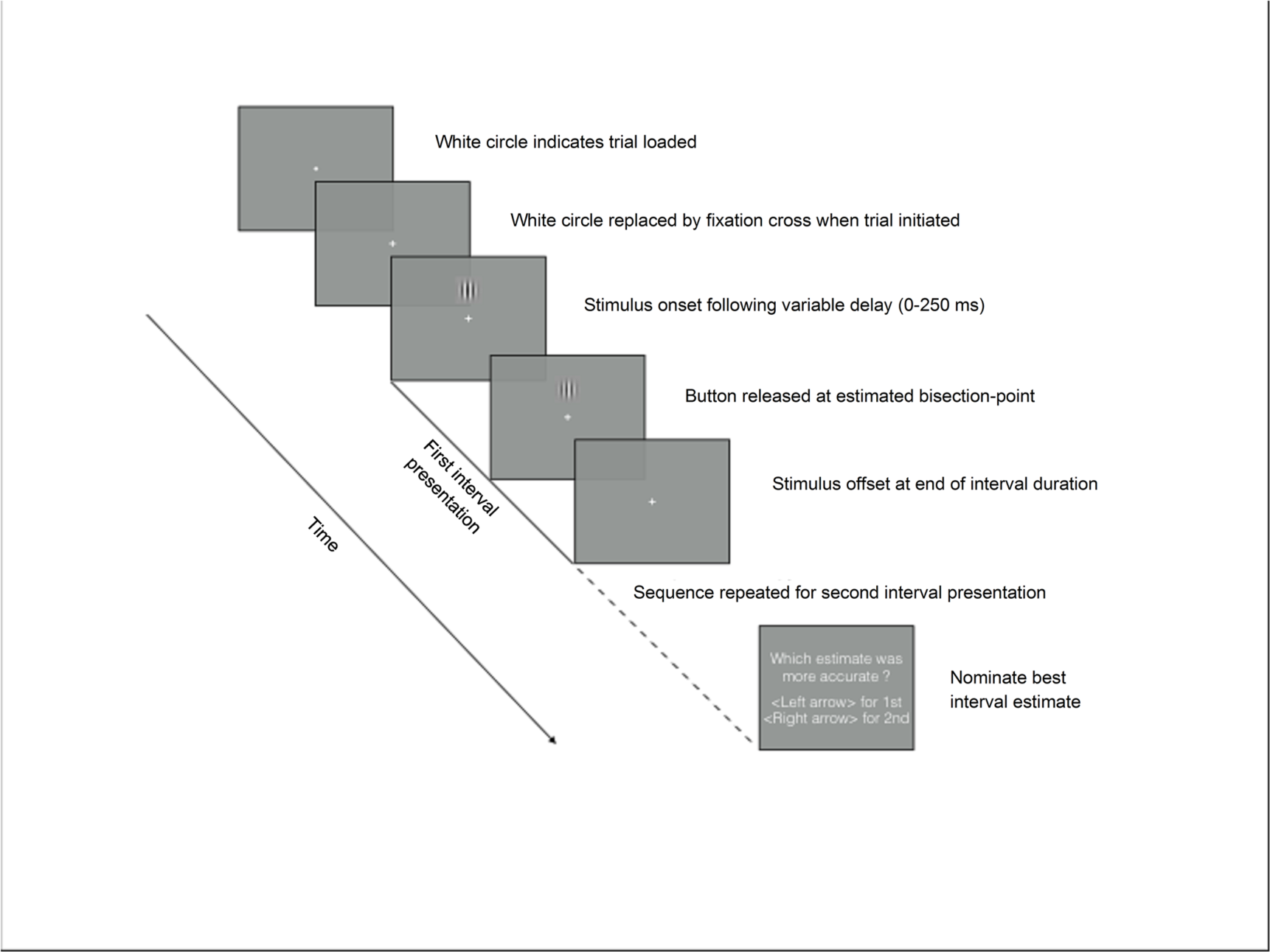
Schematic illustration of the modified temporal-bisection task trial. The stimulus is present for the full duration regardless of the estimated bisection-point. Auditory feedback can be provided on whether the participant had correctly chosen the interval closest to the actual mid-point after the second sequence.

### Bisection

Traditionally, in a standard temporal bisection paradigm, participants are given two ‘standard’ or reference intervals, one shorter and one longer, which they are required to remember. They then judge a series of single ‘test’ intervals to be closer to the shorter or to the longer standard interval, effectively bisecting the perceived difference between the two reference intervals: the test interval that is judged to be closer to either standard half the time is the perceived bisection point (Kopec & Brody, 2010). For instance, if the shorter reference was 1 second and the longer standard was 3 seconds, the inter-reference interval is 2 seconds and the veridical bisection point is also 2 seconds. The data collected from this paradigm allows construction of a psychometric function to identify the perceived midpoint in the interval between the two standards and the method is akin to the standard psychophysical Method of Single Stimuli (Macmillan & Creelman, 1991) used most effectively in measuring speed perception (McKee et al., 1986).

### Reproduction

Reproduction has, perhaps more simply, presented the participant with an interval, demarcated through whatever means (a continuous tone or light, or a pulse at the beginning and end of the interval, is common) and then asked the participant to reproduce that interval through some means such as a button-press or finger-tap. The result is then expressed as a mean and variance in the reproduced interval compared to the test duration.

We have developed a combination of these two approaches which provides a behavioural measure that allows us to observe the participants’ estimate of perceived duration in a way that is not possible in the case of the perceptual comparisons generally used in the literature (Grondin, 2010). This paradigm also affords greater flexibility with the collected data, providing both a trial-by-trial estimate of a given test duration and a cumulative summary of the mean and variance of the estimate. Importantly, we also incorporate a metacognitive component whereby the participant is required to examine their own recent performance.

### Metacognition of time

Metacognition is defined as awareness and understanding of one’s own thought processes (Fleming & Frith, 2014; Terrace & Metcalfe, 2005), and we suggest that the awareness of time-passed is crucial for interaction with the external world and the ongoing events defining that world (Franklin et al., 2020; Zacks, 2020). In the current context, we refer to the awareness of one’s own performance in the temporal task, with the intention of obtaining an objective measure of a participant’s confidence in their performance (Caziot & Mamassian, 2021). We are interested in the potential to measure the (raw) ability of participants to bisect an interval and to also determine how well participants believed they performed in the task. Given that the percept of time is a fundamentally subjective estimate of the passage of physical time in the external world, which can be so easily disrupted by many factors both commonplace and otherwise (Ayhan et al., 2012; Brown & West, 1990; Cicchini & Morrone, 2009; Droit-Volet & Meck, 2007; Grommet et al., 2011; Maarseveen et al., 2018; Morgan et al., 2008), measuring metacognition in temporal tasks is an important extension to the current standards. We introduce this possibility here by asking the subjects to compare two consecutive time interval estimates and decide which they thought was the closer estimate to the actual midpoint of the test interval. Feedback can be provided on the accuracy of this second-order judgement.

### Confidence

Subjective confidence in a decision judgement can be argued to be an outcome of the metacognition related to that decision; this relationship could then provide a window to the underlying mechanisms of a given decision (reviewed in Mamassian, 2016). There have always been issues with measuring confidence in a judgement since it is, by definition, a subjective measure that is notoriously hard to standardize across observers. The commonplace methodology of using a Likert scale to rate the degree of confidence in a given judgement is fraught with error (Caziot & Mamassian, 2021; Mamassian, 2016; Morgan et al., 1997). We suggest that the forced-choice measurement introduced in this paradigm overcomes the problems associated with Likert scale confidence measures (such as variable use of the full range between individuals, the underlying assumption of linearity across the range and variability both within and between subjects) and, we argue, provides an objective and cumulative measure of confidence on a trial-by-trial basis. Furthermore, the unique structure of the data collected on the first- and second-order judgements described here allows a genuine ideal observer analysis of the data from each participant and each condition. Our approach is substantially similar to that of Caziot and Mamassian (2021) in the current special issue, who argue that confidence should be considered a measure of internal consistency rather than external veridicality.

Here we examine bisection performance in subsecond and suprasecond intervals coded by visual, audio or audiovisual stimuli and to see whether stimuli with two coherent sources of information (audiovisual) improves performance, and consistency in the knowledge of that performance, over that with a single source alone (visual or audio). We then model both aspects of that data, bisection and metacognition, using a Bayesian approach that allows comparison to an ideal observer.

## Method

### Participants

Participants (aged 23 to 53) were recruited through convenience sampling (N= 5; 3 of them were females). No one reported any history of neurological, psychiatric disorder or were taking ongoing medication. All participants had normal (or corrected-to-normal) vision and hearing. Experiments were approved by the Human Ethics Advisory Group at the University of Melbourne. Participants provided informed consent for participation and academic use of their (anonymous) data. Participants’ names were coded as: JCZ, RXL, SC, SLY, WLX and were reimbursed for their time ($15/hour) with the exception of WLX and SC (authors).

### Apparatus and Stimuli

Stimuli were developed using the Psychophysics Toolbox, Version 3 (Brainard, 1997) and MATLAB R2018a software package (The Mathworks Inc, Natick, MA). Stimuli were either visual (V), auditory (A) or a synchronised combination of the two (AV). A Mac Pro (early 2009, OS X El Capitan) computer was used to run the software while time-critical interval-judgement responses were collected via a calibrated Cedrus RB-530 (Cedrus Corporation, San Pedro, CA) response pad and metacognitive judgements via a Macintosh keyboard.

The visual stimulus was displayed on a SONY Trinitron Multiscan G520 monitor (resolution = 1600 × 1200 pixels, frame rate = 100 Hz, mean luminance = 40 cd/m^2^, CIE co-ordinates {x = 0.333, y = 0.377}) within a stationary circular envelope (diameter = 4 °), the lower edge being located 4 ° above a central fixation spot. The audio was delivered through Sennheiser HD 25-mk2 headphones and a Roland UA-M10 external USB DAC.

The visual stimulus consisted of a vertically orientated sinusoidal luminance grating (spatial frequency = 1 cycle/°; 0.8 Michelson contrast) presented against a grey background and the mean luminance of the display The auditory stimulus was a 480 Hz pure tone set a at moderate volume around 10 times detection threshold. The synchronised stimulus was the combination of both audio and visual stimuli from onset to offset. Both stimuli were presented in rectangular temporal envelopes as onset and offset were critical aspects of the stimuli and any temporal smoothing would affect their temporal definition. We are aware of the effect of any temporal artefacts introduced by this kind of envelope.

### Procedure

The experiment comprised the three stimulus conditions presented according to the testing schedule summarised in Table 1. Three sessions were carried out over three weeks. All sessions were completed in the same order for each participant to account for and order effects when comparing between subjects. Each session took approximately 3 hours to complete, with a break of 15 minutes between the 1500ms and the 3000ms condition. The testing schedule is summarised in Table 1.

**Table 1:** V = the visual stimulus condition; A = the auditory stimulus condition; AV = the audio-visual stimulus condition.

| Session numbers | Stimulus modality | Duration condition<br>(ms) |
| --- | --- | --- |
| Session 1 | V | 1500 |
|  |  | 3000 |
| Session 2 | A | 1500 |
|  |  | 3000 |
| Session 3 | AV | 1500 |
|  |  | 3000 |

**Table 1.** Mean (and 95% HDI’s) for subject level variance parameters and subject correlation parameter. Subject level parameters for each condition

| 1500 msec Duration Condition |  |  |  |  |  |  |  |  |  |  |
| --- | --- | --- | --- | --- | --- | --- | --- | --- | --- | --- |
| Subject | Interval 1 |  |  | Interval 2 |  |  | Metacognitive |  |  |  |
| | V | A | AV | V | A | AV | V | A | AV | $\rho$ |
| 1 | 0.15 (0.14, 0.16) | 0.14 (0.13, 0.15) | 0.13 (0.12, 0.14) | 0.13 (0.12, 0.14) | 0.17 (0.16, 0.18) | 0.10 (0.10, 0.11) | 0.40 (0.30, 0.64) | 0.48 (0.35, 0.84) | 0.31 (0.22, 0.57) | 0.47 (0.02, 0.96) |
| 2 | 0.14 (0.14, 0.15) | 0.20 (0.18, 0.21) | 0.20 (0.18, 0.21) | 0.14 (0.13, 0.15) | 0.22 (0.21, 0.24) | 0.20 (0.19, 0.21) | 0.66 (0.42, 1.45) | 0.71 (0.45, 1.57) | 1.13 (0.62, 3.67) | 0.58 (0.04, 0.98) |
| 3 | 0.30 (0.28, 0.32) | 0.29 (0.27, 0.31) | 0.21 (0.20, 0.23) | 0.19 (0.18, 0.21) | 0.29 (0.28, 0.31) | 0.26 (0.24, 0.28) | 0.29 (0.24, 0.35) | 1.30 (0.82, 2.99) | 3.00 (1.76, 8.19) | 0.29 (0.02, 0.70) |
| 4 | 0.26 (0.25, 0.28) | 0.27 (0.26, 0.29) | 0.17 (0.16, 0.18) | 0.30 (0.28, 0.32) | 0.25 (0.24, 0.27) | 0.16 (0.15, 0.17) | 0.89 (0.59, 1.89) | 0.64 (0.43, 1.39) | 0.98 (0.53, 3.37) | 0.58 (0.05, 0.98) |
| 5 | 0.12 (0.11, 0.13) | 0.15 (0.14, 0.16) | 0.10 (0.09, 0.10) | 0.14 (0.13, 0.15) | 0.15 (0.14, 0.16) | 0.11 (0.10, 0.11) | 0.57 (0.38, 1.10) | 0.39 (0.28, 0.68) | 1.05 (0.61, 2.97) | 0.60 (0.04, 0.99) |
| 3000 msec Duration Condition |  |  |  |  |  |  |  |  |  |  |
| Subject | Interval 1 |  |  | Interval 2 |  |  | Metacognitive |  |  |  |
| | V | A | AV | V | A | AV | V | A | AV | $\rho$ |
| 1 | 0.30 (0.29, 0.32) | 0.20 (0.19, 0.22) | 0.32 (0.31, 0.35) | 0.32 (0.31, 0.34) | 0.22 (0.20, 0.23) | 0.24 (0.23, 0.26) | 1.39 (0.95, 2.54) | 1.04 (0.70, 1.97) | 0.51 (0.41, 0.69) | 0.60 (0.05, 0.98) |
| 2 | 0.24 (0.23, 0.26) | 0.21 (0.20, 0.22) | 0.23 (0.22, 0.25) | 0.25 (0.24, 0.27) | 0.19 (0.18, 0.20) | 0.25 (0.24, 0.27) | 1.26 (0.83, 2.47) | 1.23 (0.80, 2.50) | 0.55 (0.39, 0.93) | 0.62 (0.05, 0.99) |
| 3 | 0.40 (0.38, 0.43) | 0.59 (0.55, 0.63) | 0.54 (0.51, 0.58) | 0.37 (0.35, 0.40) | 0.51 (0.48, 0.55) | 0.39 (0.36, 0.41) | 2.25 (1.49, 4.25) | 1.93 (1.40, 3.14) | 0.96 (0.76, 1.32) | 0.80 (0.42, 0.99) |
| 4 | 0.25 (0.23, 0.26) | 0.23 (0.22, 0.25) | 0.20 (0.19, 0.22) | 0.26 (0.24, 0.27) | 0.30 (0.28, 0.32) | 0.19 (0.17, 0.20) | 0.78 (0.56, 1.25) | 0.60 (0.46, 0.88) | 0.35 (0.27, 0.49) | 0.68 (0.07, 0.99) |
| 5 | 0.24 (0.23, 0.26) | 0.15 (0.14, 0.16) | 0.11 (0.10, 0.11) | 0.26 (0.24, 0.28) | 0.16 (0.15, 0.17) | 0.12 (0.12, 0.13) | 0.90 (0.62, 1.73) | 0.44 (0.30, 0.86) | 0.61 (0.40, 1.26) | 0.32 (0.01, 0.80) |

In each experimental session, participants were tested with one stimulus condition (the visual, auditory or audio-visual stimulus), which comprised 500 trials presented in blocks of 100 trials. Each stimulus condition was tested at two durations (1500 ms (bisection of 750ms), 3000 ms (bisection of 1500ms)).

### The modified temporal-bisection paradigm

Each trial involved a temporal estimation phase wherein participants were required to make two interval-estimates immediately followed by a metacognitive appraisal where participants were required to identify which of the two precedings interval-estimates was, in their opinion, closest to the predefined target duration (the bisection point). A schematic representation of the generic trial structure is depicted in Figure 1 for a visual example. The stimulus can be of any modality as long as it can be bisected in time. In the data discussed here, the stimulus was always presented for the duration to be bisected but we have also used two pulses, one at the beginning and one at the end of the interval, as a modification of the single continuous stimulus, and varied the modality of the stimulus between the intervals to be compared (Cropper et al., 2018).

All trials consisted of two identical intervals (one of which is represented in Figure 1), each being demarcated through the continuous presentation of the test stimulus; in the example illustrated in Fig 1, the temporal cue was solely visual. The participant was told the interval duration that was to be used for the entire block of trials and that they were required to estimate half of this interval, bisecting the time period that the stimulus was present on the screen. The first interval was initiated when the participant depressed the response pad button. To avoid the use of rhythm in the task (which we found, through pilot testing, made the task trivial), a random delay, ranging from 0 to 250 ms, was inserted after the button press and prior to the appearance of the stimulus. Upon appearance of the stimulus, the participant maintained button depression until the perceived interval midpoint, when they released the button. The stimulus remained on-screen for the full duration of the interval. This procedure was then repeated for the second interval estimate following a 500ms inter-stimulus interval.

After making a pair of bisection-point estimates, participants were prompted to make a two-alternative forced-choice response via the keyboard arrow keys to indicate which of their estimates they deemed closest to the target duration; i.e., their ‘best’ estimate. This retrospective judgement constitutes the temporal metacognition component of the task. This paradigm therefore contains a first-order judgment, which is a combination of temporal-bisection and interval-reproduction, followed by a second-order forced-choice judgment of one’s own accuracy (Caziot & Mamassian, 2021; Mamassian, 2016).

### Treatment and analysis of the data

One of the particular benefits of the modified bisection paradigm is that the data collected can be examined both as a trial-by-trial analysis of the subject performance and, more traditionally, as an expression of performance over the whole experimental condition.

### Basic raw data and sorting of individual responses

Example raw data for one observer (AK) is shown in Figure 2 as frequency histograms plotting the estimated half-point of the 1500-ms-interval for 500 trials. The data is paired into three ways of presentation sorted by rows on the figure. The first row plots, as a histogram, the bisection data for the first and second interval estimates respectively. The second row re-sorts the data into the perceived best (left-hand side) and perceived worst of each pair. The third row plots the actual best and worst estimates, i.e. the ideal observer response. Each of the distributions has a mean and variance that can be used to summarise performance across all the trials and the trial-by-trial data can be analysed as a time-series. The individual trial estimates, followed by each pairwise analysis of performance, allow a subjective ideal Observer analysis for each subject and each set of conditions. The bimodal pattern of the ‘ideal worst’ response is exaggerated in this example (real) data set but is a direct and predictable product of the data-sorting algorithm and depends upon the trial-by-trial values of the particular data set which are influenced by both subject & condition. The ‘ideal worst’ response will always have the estimate further from the mean of any given pair creating a distribution containing the tails of the original population, the ‘ideal best’ response set receiving the data closest to the mean. This allows the difference in variance between the best and worst distributions to be used as a measure of the knowledge of the observer of their own performance but also potentially changes the shape of the distribution (Corcoran et al., 2018).

**Figure 2.**
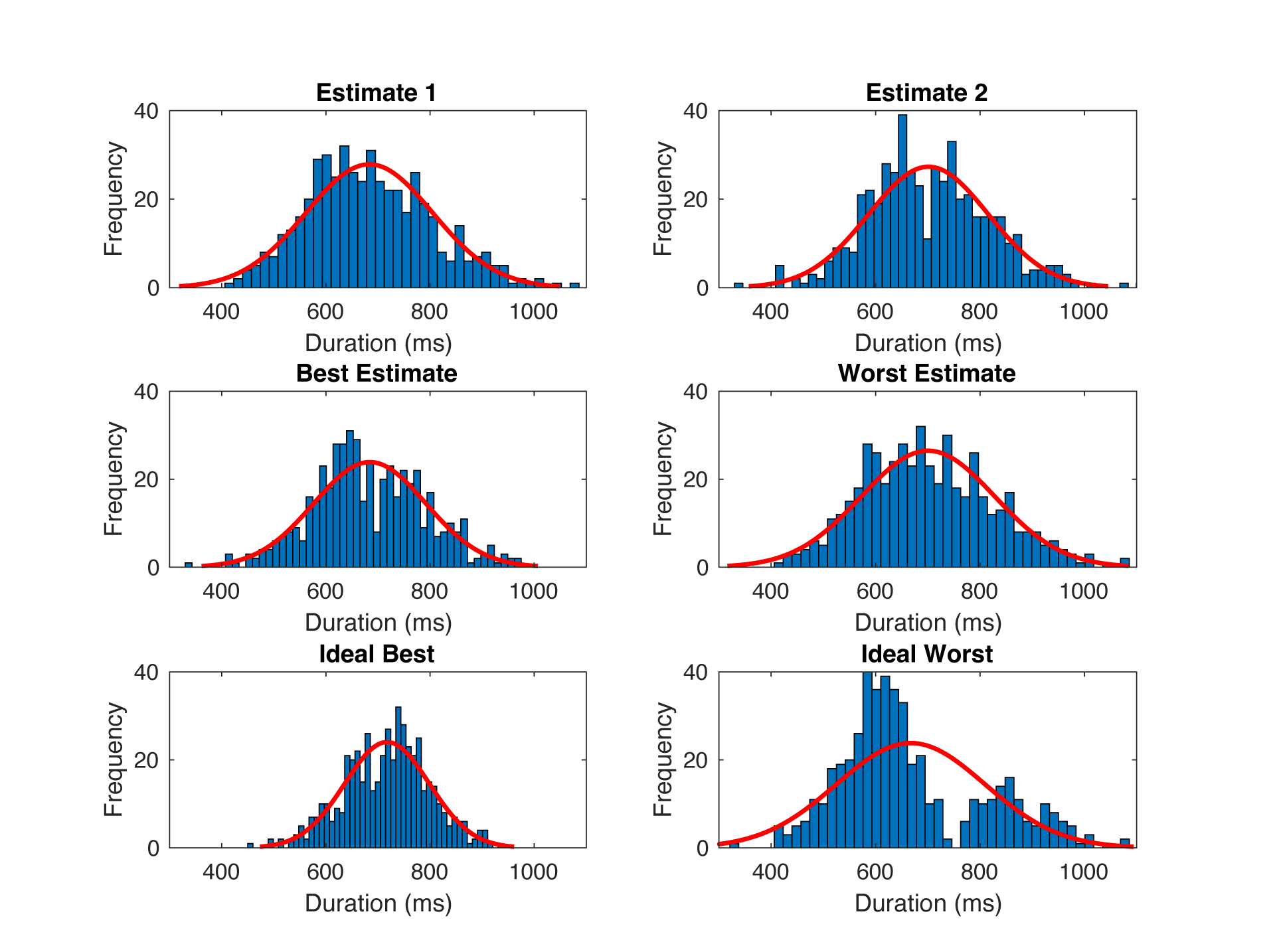
Histogram plots using 50 bins from estimates of zero, to the total stimulus duration for one observer (AK) in the visual same-modality 1500 ms stimulus duration condition. Red lines indicate Gaussian of best fit. To characterise observers’ second-order responding, the trial-by-trial data was redistributed according to observers’ best estimate judgements, to gain mean and variance values of subjective best estimates, and by complement, subjective worst estimates, across trials within a condition. The data was also redistributed according to the ideal best, and ideal worst estimates, to gain the mean and variance in the case of ideal observer performance (ie. the estimate distributions expected if observers had perfect insight into their own performance).

### Analysis of performance over the duration of the experiment

Figure 3 plots the performance on a trial by trial basis for a single interval of the pair (Interval 1 in this case) over the entire 500 trials. The individual bisection estimate (y-axis) is plotted against the trial number (x-axis). The binned means (bins of 50 trials, data points in middle of bin), with 95% confidence intervals superimposed on the trial-by-trial data. The benefit of looking at the data this way is that one can see how performance changes over the duration of the experiment.

**Figure 3.**
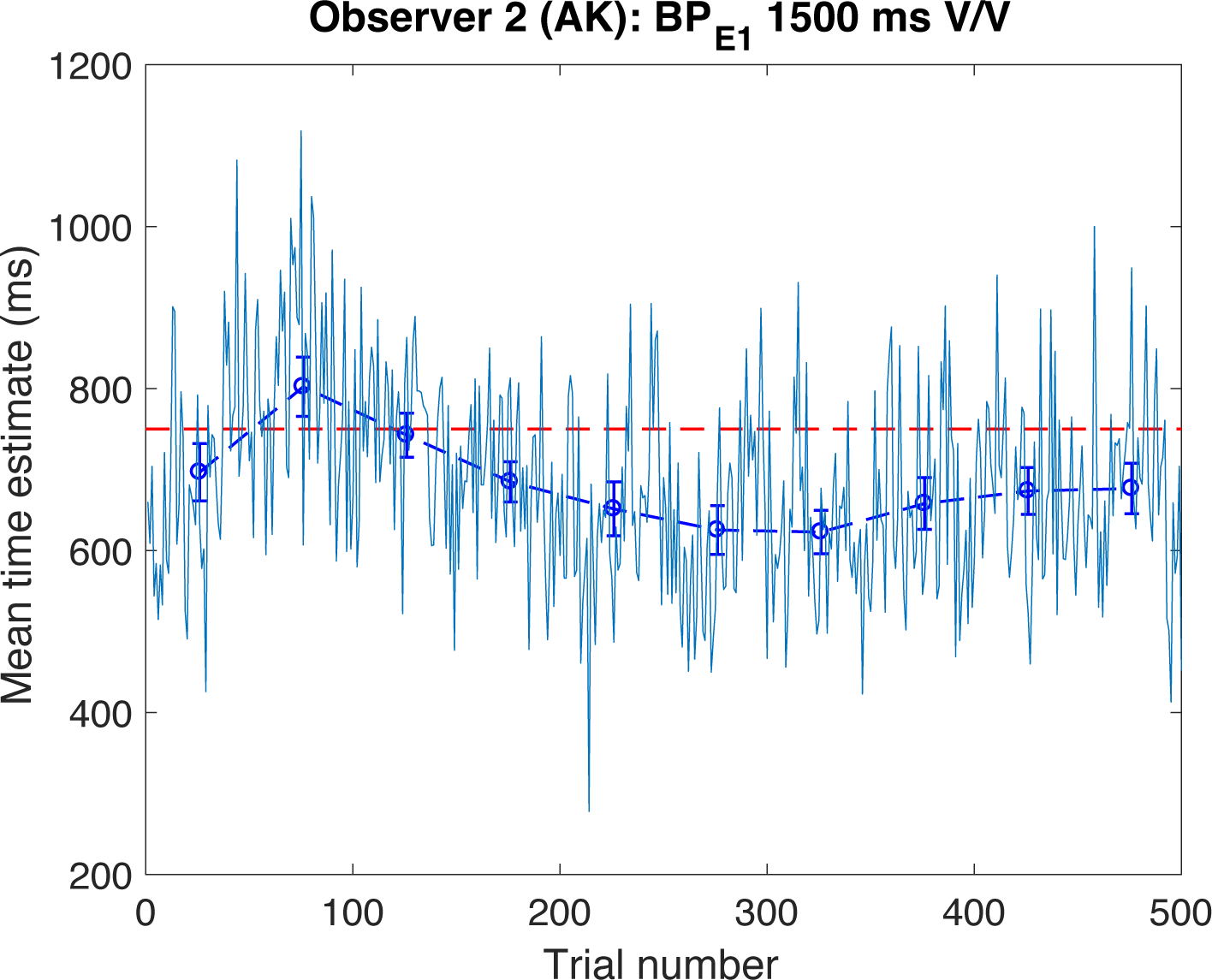
Raw trial-series of BPE1 for observer 2 (AK; teal dashed line), for the visual same-modality (V/V) 1500ms stimulus duration condition. Note that BP_A_ = 750ms is shown as a horizontal red dashed line. Binned means with 95% CI shown in royal blue, connected by dashed lines.

This trial-by-trial approach to the analysis can be extended to examine how previous trial performance affects the current trial either globally by curve-fitting the data or more locally by conducting an autocorrelation of the data. Each of these approaches offers some possibility of examining how the internal representation of the target interval builds up for the participant over the period and how recent performance affects the current decision.

### Summary analysis: re-Sorted data

As implied in Figure 2, a summary analysis of performance is perhaps the most obvious and common way of looking at the data. The mean and variance of the two intervals (Row 1 of Figure 2) give the accuracy and precision respectively of the two bisection estimates and are the standard way of representing this kind of data, providing a means of comparing performance in this paradigm to others in the literature. This approach also allows the coefficient of the variation (i.e. standard deviation of estimated duration / mean estimated duration) to be used to express the adherence, or otherwise, to Weber’s law, which predicts a constant coefficient across different bisection periods and is an enduring critical component of the scalar expectancy theory of time (Corcoran et al., 2018; Grondin, 2014).

#### Metacognition, Feedback and Ideal Observer Analysis

Re-sorting the individual interval 1 & 2 trials into perceived best and perceived worst and ideal best and worst gives some insight into how effectively the individual subjects in each condition are able to access and reflect upon their own performance during the task. We expect (and observe (Corcoran et al., 2018; Cropper et al., 2018)) that the difference between variance of the estimates will be minimal in the Interval 1/2 condition, and maximal between ideal best/worst. The difference in the subjective best/worst case will be somewhere between these two extremes and this difference can be used to quantify the degree of knowledge each subject had of their own performance in the bisection/reproduction task.

One way of summarizing this is as follows:

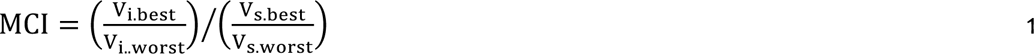

Where the MCI is the MetaCognitive Index for a given subject and condition, V is the variance of the ideal or subjective best/worst interval (indicated by the subscript). If the subject has little knowledge of their own performance, then the variance of the ‘best’ and ‘worst’ intervals will be substantially the same and the ‘subjective’ denominator of Eq 1 (V_s.worst_) will tend toward a value of 1 and the overall MCI will be the ideal numerator; tending to, but not reaching, zero. If the subject has perfect (ideal) knowledge of their own performance and then both (ideal) numerator (V_i.best_)and (subjective) denominator of Eq 1 will be the same, giving a result of 1.

To account for the bimodal distribution in the ideal worst data set in Figure 1, and not artificially inflate the value of the ideal condition, the mean of the ideal best and ideal worst variance may be used:

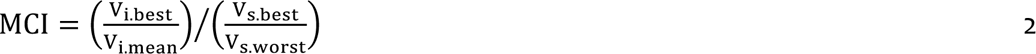

Where V_i.mean_ is the (V_i.best_ + V_i.worst_)/2.

MCI values calculated for the example data in Figure 1 are 0.47 (Eq 1) or 0.69 (Eq 2). This better estimate (Equation 2) of subjective appraisal is consistent with a more conservative estimate of ideal worst variance.

### Empirical Results

Subjective ‘best’ estimates for the bisection of the two presentation intervals (1500ms and 3000ms) are presented and bar-charts in Figure 4 for the 5 observers across each of the three conditions (Visual, Audio and Audiovisual). There is some individual difference in performance, as expected (Corcoran et al., 2018), but all subjects do relatively well in the task in all conditions and also show a good degree of consistency with small error bars (+/-SEM). Overall performance appeared closer to the veridical bisection point in the longer duration condition. Figure 5 replots this data as the bisection point in each condition relative to the total duration which tends to reduce this apparent difference between durations but emphasises the individual differences in performance.

**Figure 4.**
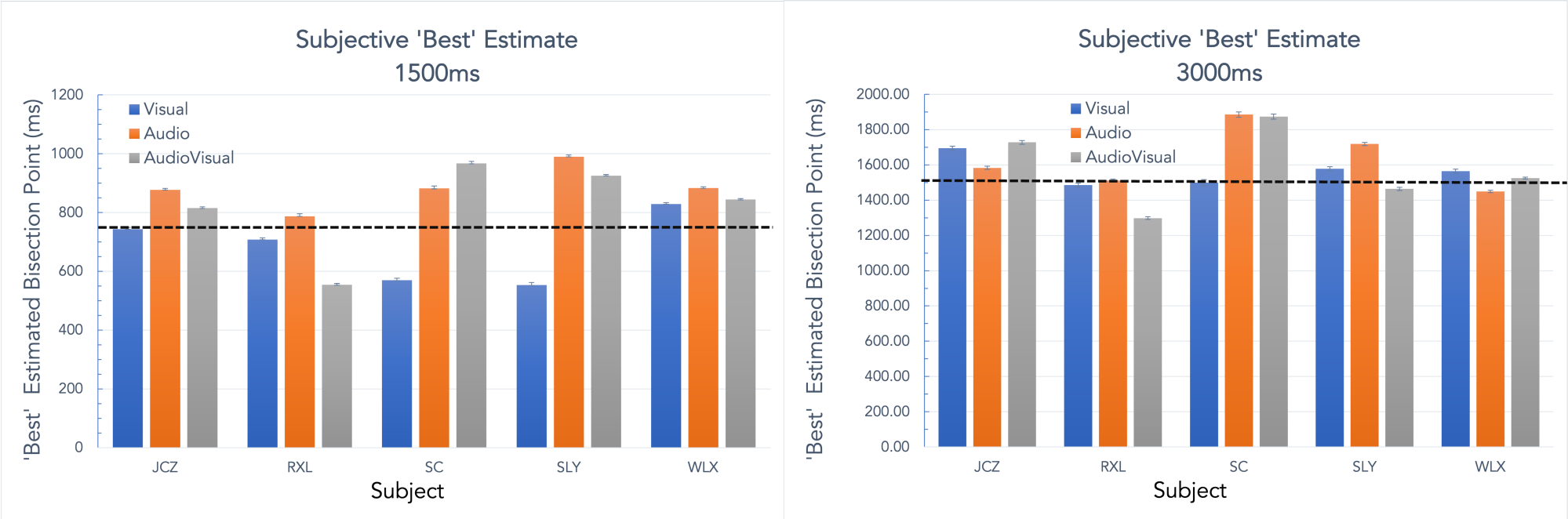
“Best” estimate of the interval bisection plotted for 5 subjects across three conditions (Visual, Audio and Audiovisual). Error bars are +/-SEM and the dotted line indicates the veridical bisection point.

**Figure 5.**
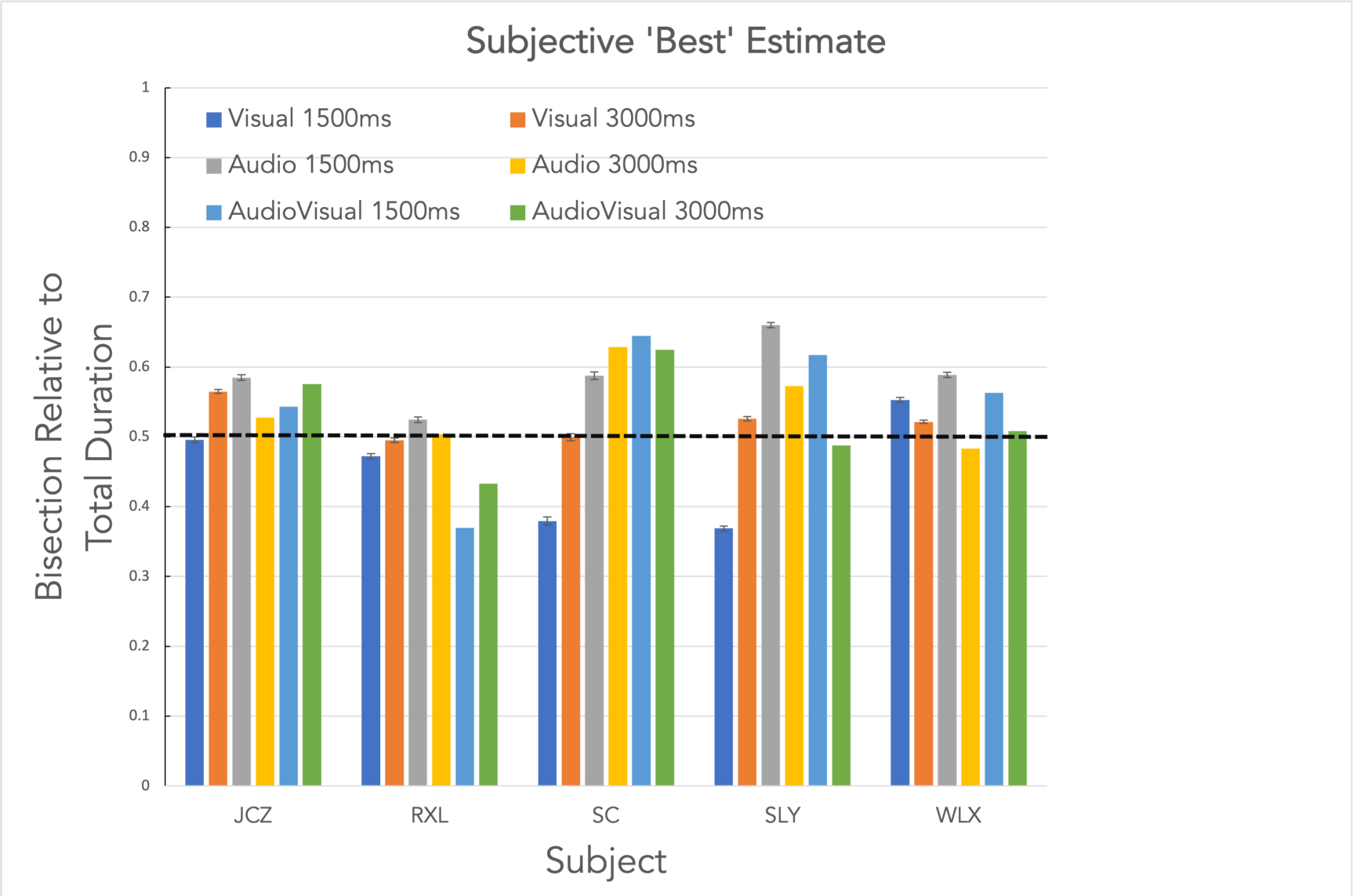
“Best” estimate of the interval bisection plotted relative to the total duration for the 5 subjects across conditions (Visual, Audio and Audiovisual, 1500 and 3000ms). Error bars are +/-SEM and the dotted line indicates the veridical bisection point.

Figure 6 plots the metacognitive index (MCI) for each subject and condition shown in Figure 4. The closer the MCI value is to 1, the better the knowledge the subject has of their own performance in each trial-pair. There is a tendency for the Audiovisual condition to give a higher MCI than either modality alone in three of the subjects (RXL, SC & SLY) although not with subjects JCZ & WLX.

**Figure 6.**
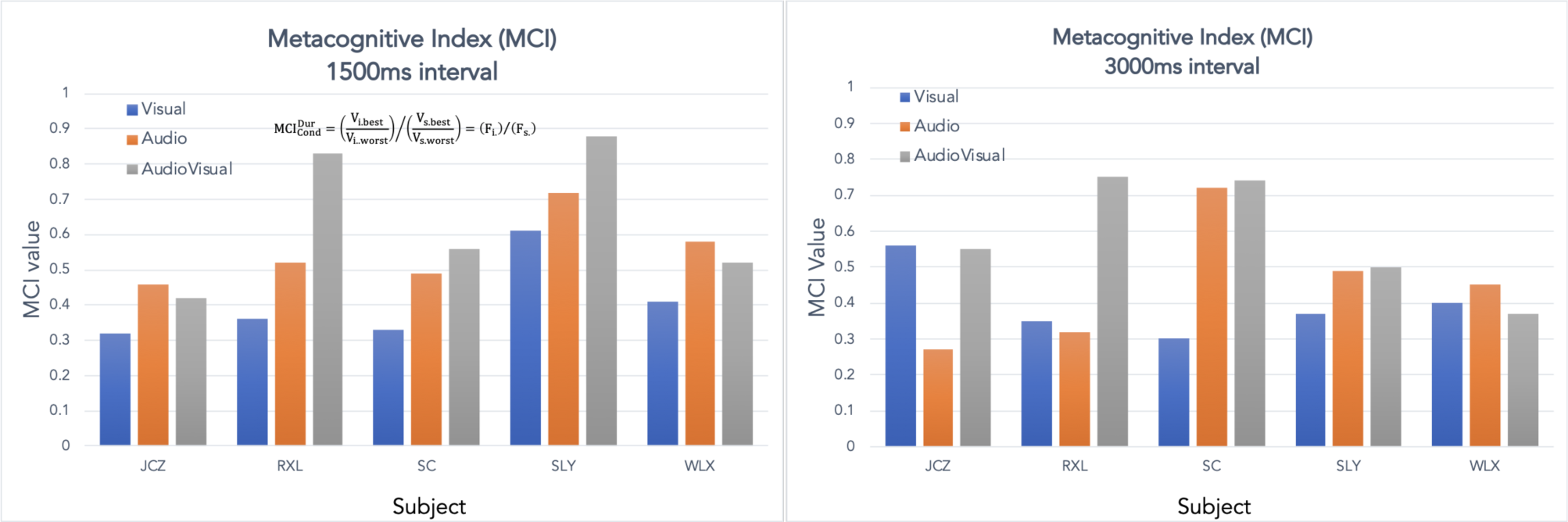
Metacognitive Index (MCI) for each subject and condition, the bisection data for which is plotted in Figure 4.

### Bayesian Model of Bisection Estimates and Metacognitive Accuracy

We developed a model of metacognitive accuracy that accounts for both bisection estimates and metacognitive decisions simultaneously and allows for comparison of a subject’s actual performance to an ideal observer in an alternative manner to the MCI defined above. The benefit of this analysis is that the parameters of the model can be estimated hierarchically, allowing each observer to inform the group-level distribution. The group-level distribution also “shrinks” estimates of subject-level parameters toward the group average (Gelman, Carlin, Stern & Rubin, 2004; Davis-Stouber, Dana & Rouder, 2018) allowing for more accurate estimates at the group level.

We modelled the metacognitive decision on each trial, *r_i_*, as a Bernoulli distribution with a parameter, *p_i_*, indicating the probability that interval 1 was chosen as the more accurate interval on trial i. This parameter was derived from a comparison of the perceived error in each interval as follows: Let Δ_1i_ and Δ_2i_ be the error in intervals 1 and 2, respectively, on trial i. That is:

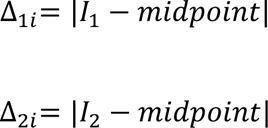

where I1 and I2 are the bisections made by the participant on intervals 1 and 2, respectively, and *midpoint* indicates the true bisection point of the interval. We model the error in the midpoint estimation by assuming that the observed interval estimates I1 and I2 are normally distribution with an observer specific standard deviation (which is unique to each condition – modality × duration – and interval).

The model assumes that the participant chooses the option with the smaller perceived error. To implement this, we find the difference between the error in each interval: *d*_Δ_ = Δ_1_ − Δ_2_. (Here, we’ve suppressed indexing by trial for simplicity). If the difference is negative, then the error in the first interval is smaller, and the participant should choose the first interval. If the difference is positive, then the error in the second interval is smaller, and the participant should choose the second interval.

We assume that the difference between the error estimates is normally distributed around the true error difference, with a standard deviation representing the accuracy of the error estimation.

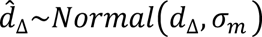

In the ideal observer model, the standard deviation would approach zero and consequently the representation of the error would approach the true error. *p*_i_, which determines the Bernoulli outcome of the metacognitive decision, then equals integral from the distribution of 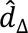 between −∞ and 0.

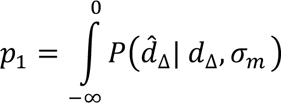

When the error in interval 1 is smaller than in interval 2, the distribution will have most of its area in the negative region, and hence, when variance is small, *p*_1_ will be close to 1 (see Figure 7). When the error in interval 2 is smaller than in interval 1, the distribution will have most of its area in the positive region and *p*_i_ will be near 0 (i.e., interval 2 will be preferred). As the standard deviation approaches 0, the model will choose the interval with less error more accurately.

**Figure 7.**
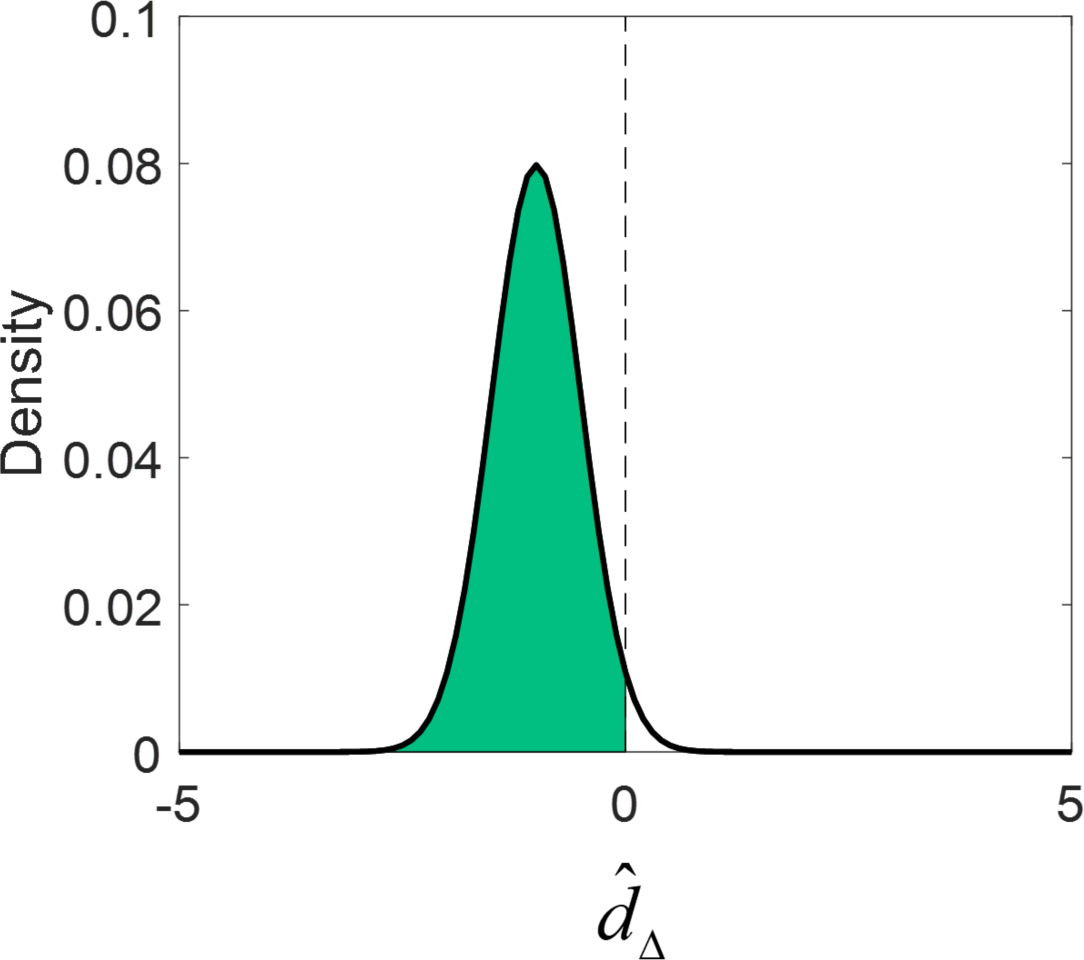
Illustration of how the probability of responding with interval 1 on a given trial is derived by integrating the distribution over the perceived difference in error.

We estimated the three standard deviations, *d*_*I*1_, *σ*_*I*2_, and *σ*_*m*_, hierarchically. We implemented the model in JAGS (Plummer, 2003) using Matlab and matjags (Steyvers, 2011) using two chains with a burn-in period of 2000 and a sampling period of 5000 samples, thinning every 20^th^ sample. We assumed a group-level distribution over the standard deviations for the V, A, and AV conditions. JAGS specifies the spread of the normal distribution in terms of precision (i.e., 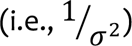); the prior distributions for each precision parameter were uniform from 0 to 100. To capture the repeated measures design, we assumed that the subject-level precision parameters were sampled from a multivariate normal distribution with a shared variance-covariance parameter and a subject-specific correlation parameter that captured how correlated the variance estimates were across the V, A, and AV conditions. All precision estimates were transformed to standard deviations for ease of interpretation.

In a nutshell, group-level standard deviation estimates for the V, A, and AV conditions measure the accuracy of the interval estimation and the metacognitive accuracy. Values near 0 indicate more accurate (veridical) estimates. At the group-level, these parameter estimates allow inference as to the difference between each modality and the combined AV condition. The subject-level parameters indicate the performance of different individuals and allow an assessment of individual differences in bisection and metacognitive judgment accuracy.

Figure 8 shows the graphical model used to estimate parameter posteriors in each duration condition. All chains showed good convergence (maximum R-hat = 1.0002; Gelman et al., 2004). Figure 9 shows the posterior density estimates for the group level parameters. The interval estimates show roughly the same variance regardless of whether the stimulus was presented Visually, Auditorily, or as a combined Audio-Visual signal. The variance of the metacognitive judgement, however, which determines the accuracy, varied depending on whether the interval was 1500 msec or 3000 msec. In the former, the AV condition was less ideal than the A or V conditions, but this pattern reversed in the latter condition. Table 1 shows the subject-level parameters in each condition; of key interest is the subject level correlation parameter which is estimated to be greater than 0 in all cases.

**Figure 8.**
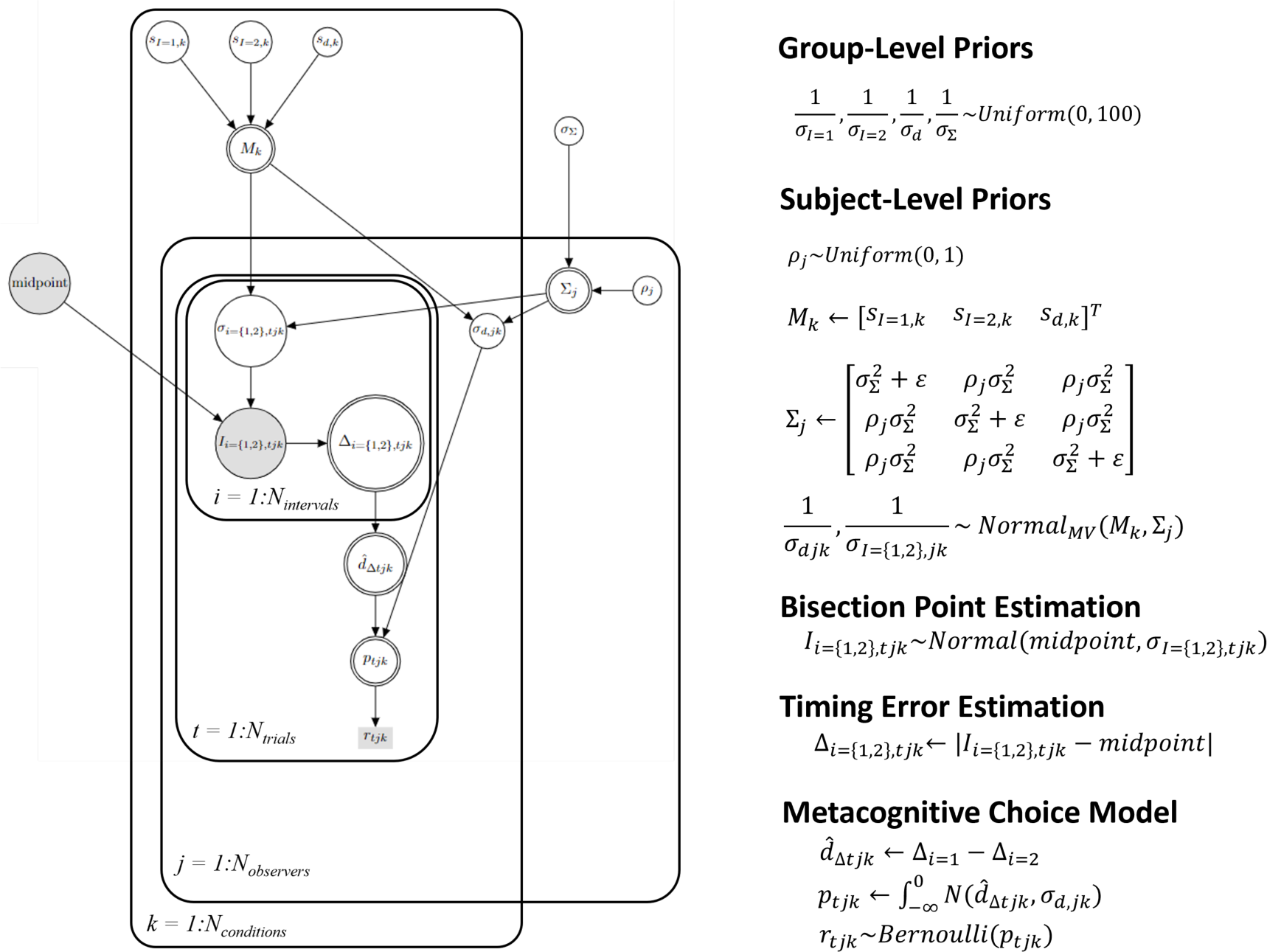
Illustration of the model as a directed graph (see e.g., Lee & Wagenmakers, 2006). Shaded boxes illustrate observed discrete variables, double circled nodes indicate derived variables, and white circles indicate continuous latent variables. The likelihood and prior distributions are listed to the right of the graphical model. Variables are defined in text; ε = .1 to ensure a positive semi-definite covariance matrix.

**Figure 9.**
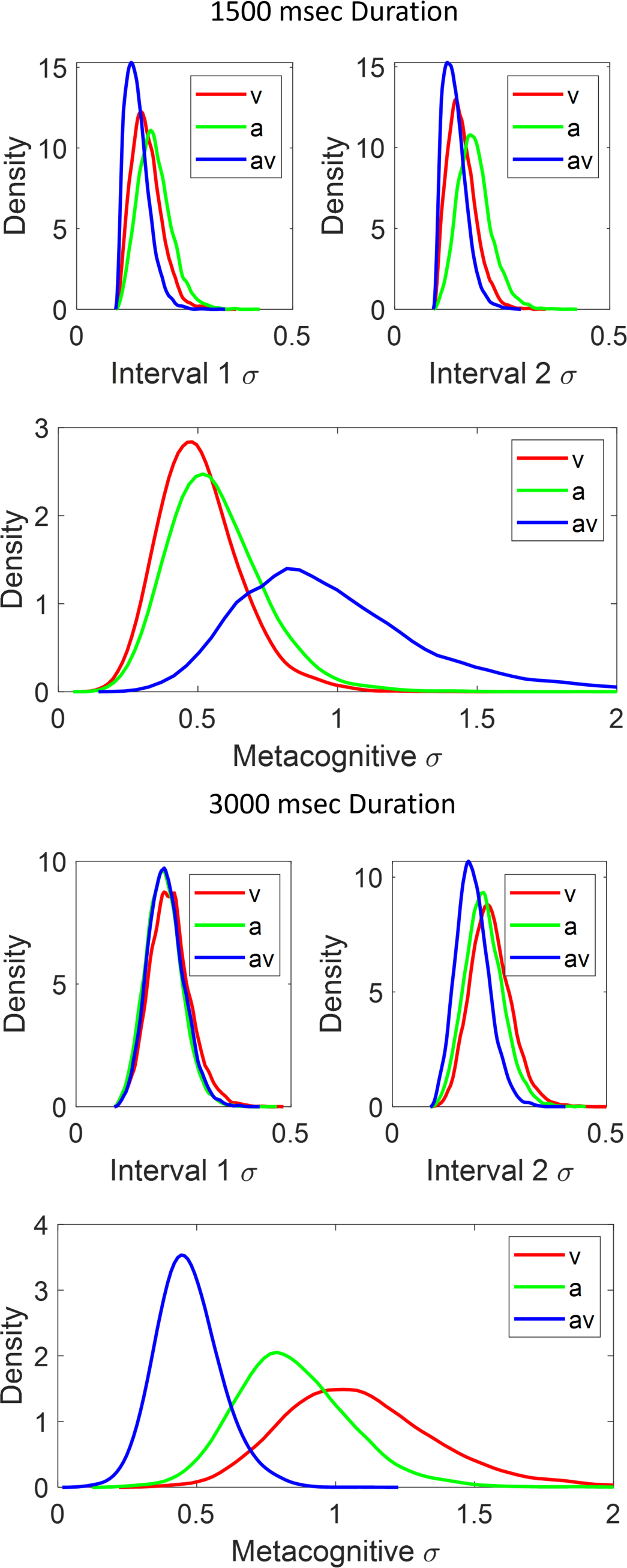
Posterior densities for the interval standard deviations and metacognitive decision standard deviations in the V, A, and AV conditions for 1500 and 3000msec durations; upper and lower panel respectively.

For the interval estimation (see Figure 9), the standard deviation estimates are all greater than 0 and vary between 0 and .5; the standard deviations, at the group level, are roughly equivalent between modality and duration conditions. This indicates that the bisection estimation was generally of equivalent accuracy between conditions. By contrast, the standard deviation estimates of the metacognitive accuracy vary not only between different modality conditions but also by duration. For the shorter duration, the AV condition metacognitive estimates are generated with greater variability than the A or V conditions, which are roughly equated. For the longer duration, the AV condition metacognitive estimates are less variable than either of the single modality conditions suggesting that the increased duration allows the dual modality condition to improve metacognitive performance consistency.

Figure 10 plots the subject-level midpoint estimates for each condition. Note that these estimates qualitatively follow the same pattern as the empirical averages shown in Figure 4 but are now additionally constrained by the group level estimates as well as the data from both intervals, not just the subjective ‘best’ interval. Finally, the group-level midpoint estimates are shown in Figure 11. In both duration conditions, the overall mean (across modalities) tends slightly toward overestimation of the bisection interval). However, this is more consistent in the longer duration condition. In the shorter duration condition, the V modality condition tends toward underestimation.

**Figure 10:**
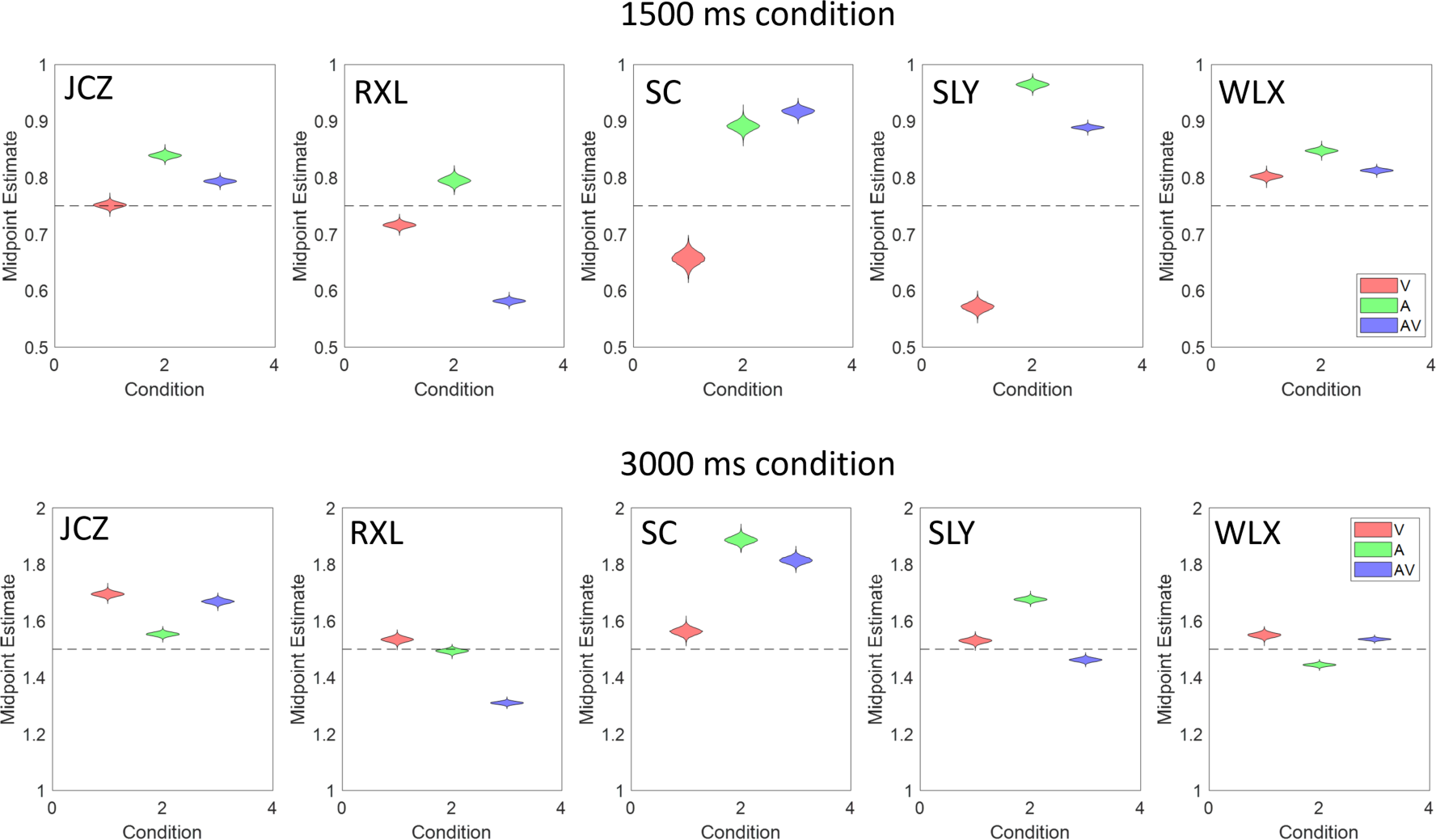
Subject-level midpoint estimates for each condition and interval duration.

**Figure 11:**
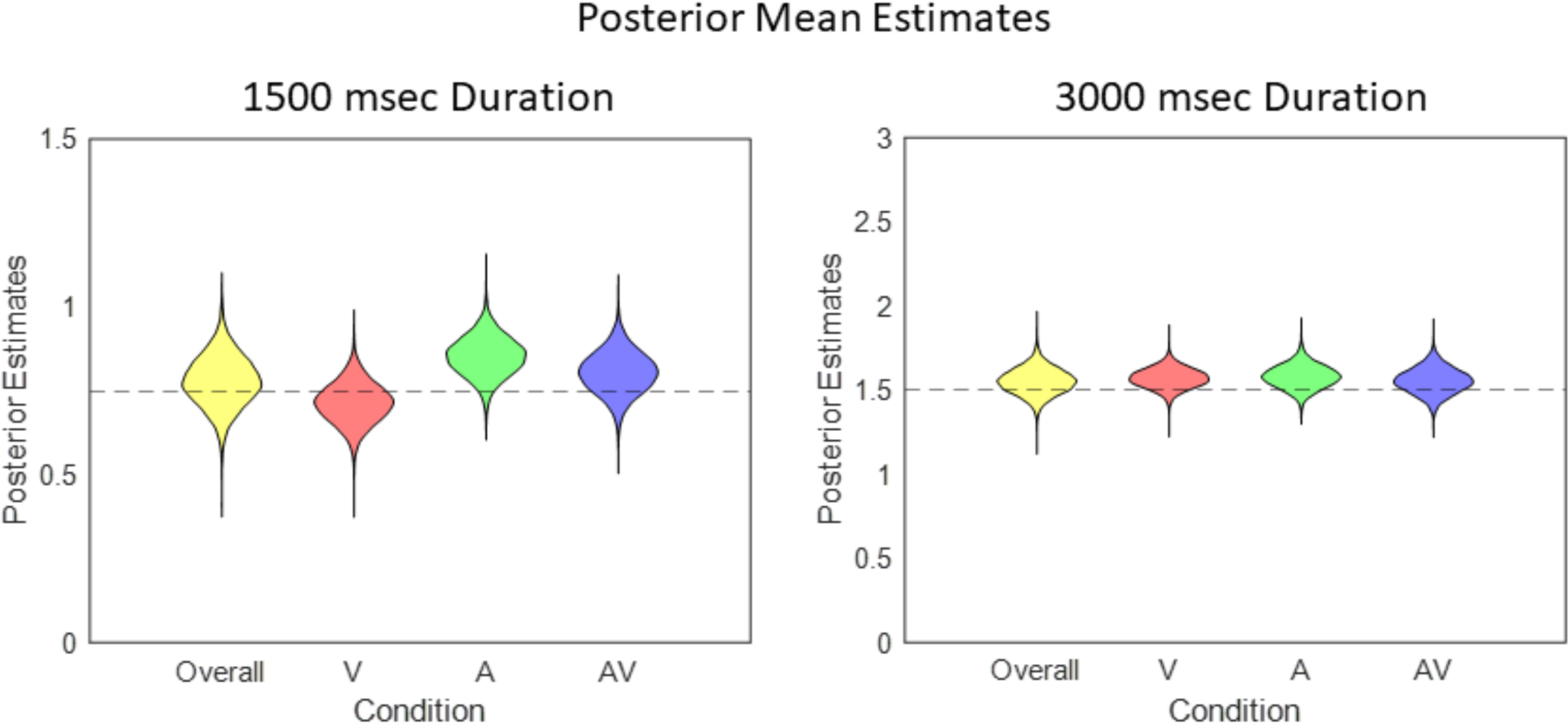
Group posterior midpoint estimates in the two interval conditions for each stimulus type and the overall mean.

## Discussion

The principal aim of this paper is to describe a novel paradigm to measure the perception and metaperception of time and suggest why we believe it is a useful contribution to the area. We also present a simple model to explain the collected data over a modest range of stimulus properties and intervals. The paradigm we describe combines temporal interval *bisection* and *(re)production* as described in the existing literature (Grondin, 2010) with an additional forced-choice measure of confidence (Mamassian, 2016). In our paradigm, we ask participants to bisect (i.e. identify the mid-point of) a temporal interval as they perceive and monitor that interval, repeat the procedure and then judge how well they performed in that pair of test-intervals. This requires the subject to build up and maintain an internal representation of the target interval that they attempt to match each time. In the second-order judgement of trial-by-trial performance, where a single trial is a pair of test-intervals, it is also possible to introduce feedback to indicate whether the interval they chose to be their best estimate was, in fact, closest to the veridical midpoint of the pair. We reiterate that this feedback (if used) does not give any information as to the accuracy of the estimate other than whether their choice was the better guess. This direct (first-order), ongoing behavioural estimate of duration followed by the second-order assessment of performance, is we believe, unique in the timing literature and offers several useful and transparent ways of analysing the collected data.

The limited data presented here and the modelling of that data suggest that while the measurement of the bisection point using this paradigm has fundamentally similar properties at both sub- and suprasecond durations, and across single and dual modality conditions, the metacognitive measurement is significantly affected by modality and duration, Specifically, in the dual-modality condition the metacognitive response shows greater variability, and therefore less internal consistency (Caziot & Mamassian, 2021) than either modality alone in the shorter duration condition, and less variability and therefore more consistency in the longer duration. The most simple explanation for this might be that more time is required for the benefit of a dual modality to be gained, and at the shorter durations the dual input may actually act as a distraction in the task despite it coding exactly the same temporal signal. The benefit of bimodal input for sensory discrimination has been shown for audiovisual signals presented for 2,5s, in the range of our longer duration stimuli here (Koene et al., 2007). The bimodal advantage in first-order discrimination may then result in greater internal consistency in the metacognition of performance as suggested in the current data, changes in stimuli and task notwithstanding.

### The Paradigm

#### Basic summary data

The calculation of the mean and variance of the estimated duration over the whole of the experiment is the simplest way of looking at the data (Figure 2, Row 1). This approach gives a measure of both accuracy (mean) and precision (variance) in the estimate and facilitates comparison to more standard methods of bisection and reproduction. We have found that even under the constraints of the larger group-design studies the vast majority of subjects can do the task with a fairly consistent degree of precision even though their perceived mid-point may not be the veridical mid-point (Corcoran et al., 2018).

#### Trial-by-trial analysis

Rather than just using the sum of all responses over the experiment, we consider one of the major benefits of this paradigm is the ability to look at the evolution of the bisection judgment over the period of the experiment. The paradigm allows the data to be considered as a time-series of responses for a given participant and condition, and the change in performance over time mapped and quantified. We suggested above that this approach can be well expressed through curve-fitting and autocorrelation, but obviously the data can be presented as appropriate for the experiment. The benefit here is that the trial-by-trial performance of a subject offers a window into their internal representation of the interval to be bisected as they perform the task.

#### Metacognition, confidence and ideal-observer analysis

Incorporating the second-order measure of performance into the analysis allows the data to be reordered in a way that affords access to how good the subject thinks they are doing in the task (Figure 2, row 2). We argue that taking this objective (forced-choice) measure of how well the participant thinks they are doing in the task (i.e. how accurate they think they are) allows a more reliable measure of subjective confidence than the standard rating approach. This property is another unique aspect of the paradigm and the way in which the bisection estimate is made.

The ability to then reorder the individual trial data into perceived and actual best/worst performance allows a given subject’s performance on the second-order task to be expressed in the context of an ideal observer analysis (Geisler, 1989; Geisler, 2003; Geisler, 2011) and for this relationship to be broken down over the duration of the experiment. The benefit of this particular paradigm for this theoretical approach is that the trial data for the ideal ‘device’ is actual data from the subject and condition in question, rather than estimated or averaged data as is often the case. This means that the degree of ideal behaviour of the system at the level of temporal metacognition and its properties over the duration of the experiment can be more accurately judged for a given set of conditions. It is also possible that this aspect of the data might allow some reasonable discrimination between underlying models of time perception by examining how they behave over multiple consecutive samples of a given epoch and how that compares to collected data (Jazayeri & Shadlen, 2010).

The decision made by the subjects is different from a standard two-interval forced choice magnitude-discrimination task. In the standard task, two sensory quantities are directly compared with one another and the decision made about which is greater. In our task, each sensory quantity (i.e. the two independent bisection estimates) is (are) compared to an internal reference (the veridical, or perceived, midpoint of the interval which is never actually given) and the judgement made of which one is closer to the midpoint, making it a metacognitive judgement. The subject cannot make the correct decision based on which appears longer or shorter directly as they might both be too short (in which case the longer one is correct) or too long (the shorter one is correct). The subject still has to choose which of those independent bisection decisions to accept based on their judgement of the reliability of their judgements. We argue that this decision process allows us to say that we have objective evidence on accessing the information path that allows the subject to make their decision

As we suggest above, looking at the variance in this way also gives us an objective measure of observer confidence in the task and the possibility of measuring how confidence in ongoing performance changes over the duration of the experiment. This may also be considered as a way of examining internal consistency which is possibly a better way of looking at the idea of subjective confidence in a given judgement (Caziot & Mamassian, 2021).

#### Feedback

The addition of feedback at this level into the paradigm i.e. only at the second-order judgement level, allows some modulation or reflection to be introduced into the metacognition (or in Bayesian terms, a prior to be updated). The only noticeable effect of this manipulation on subjects’ performance was to decrease the number of trials at which a bisection judgement somewhat close to veridical was reached (12-15 trial pairs in most subjects (Corcoran et al., 2018; Cropper et al., 2012)). An unusual property of providing feedback at this stage in the decision process is that once the perceived bisection point is estimated relatively accurately in both intervals, they potentially become indiscriminable from one another making the provision of feedback more influential on the decision process earlier in the trial series.

#### Modelling

The benefit of the hierarchical Bayesian model is that it allows for simultaneous estimation of group- and observer-level parameters and provides an estimate of the uncertainty in those parameters. In our implementation, we jointly estimate the variability of the interval estimation and the metacognitive estimates allowing for determination of the correlation in the error between these estimates. Our model is a measurement model rather than a psychological model, using the data to return useful estimates of uncertainty parameters. Nevertheless, we model the decision process in a psychologically-meaningful way (see Figure 6) drawing on principles of signal detection (Green & Swets, 1966; Macmillan & Creelman, 1991) to provide a theoretical framework for the decision. A limitation of our approach is that we assume the psychological midpoint is static across trials (contra to Figure 3). The model could also be extended by allowing the estimates to vary functionally across trials, but we leave that for future work.

## Conclusion

The paradigm described here offers a different way to measure the percept of brief intervals of time as they are experienced by the subject. The two stages of the decision give both a first-order measure of the temporal percept and a second-order metacognitive judgement of performance. The summary measures provide some equivalance to the extensive literature on time perception, while the more unique trial-by-trial data collection affords a detailed window into the way in which the system builds up a representation of the duration to be bisected, which in turn mediates future performance.

